# Welcome to the jungle: Algal turf negatively affects recruitment of a Caribbean octocoral

**DOI:** 10.1101/2020.05.27.119404

**Authors:** Christopher D. Wells, Ángela Martínez-Quintana, Kaitlyn J. Tonra, Howard R. Lasker

**Affiliations:** Department of Geology, University at Buffalo, State University of New York, Buffalo, NY; Department of Environment and Sustainability, State University of New York at Buffalo, Buffalo, NY

**Keywords:** Cnidaria, Cox mixed effects model, epilithic algal matrix, gorgonian, herbivory, mesofaunal predators, Octocorallia, Scleractinia

## Abstract

1. Algal cover has increased and scleractinian coral cover has steadily declined over the past 40 years on Caribbean coral reefs. In contrast, octocoral abundance has increased at those sites where octocoral abundances have been monitored. The effects of algal cover on recruitment may be a key component in these patterns, as upright octocoral recruits have the potential to escape competition with algae by growing above the ubiquitous algal turfs. However, the impacts of algal turf on octocorals have not been tested.
2. We used laboratory and field recruitment experiments to examine impacts of algal turf on settlement and then survival of newly-settled octocorals. Tiles were preconditioned on a Caribbean reef, allowing algae to settle and grow. Tiles were then partitioned into three treatments: lightly scrubbed (0% turf cover), left alone (19% turf cover), or kept for 15 days in a sea table without fish or large invertebrate herbivores (50% turf cover). Planulae of the common Caribbean octocoral *Plexaura homomalla* were allowed to settle and metamorphose on the tiles for six days. Tiles were then deployed onto a reef and survival of those recruits was monitored for seven weeks. Settlers that recruited to the tiles after deployment to the reef were also monitored.
3. Laboratory recruitment rate was significantly higher in lower turf cover treatments. Field survival was significantly reduced by increased turf cover; for every 1% increase in turf cover, polyps died 1.3% faster. In a model parameterized by the observed field survival, polyps exposed to 100% turf cover had a 2% survival rate over 51 days, while polyps exposed to no turf cover had a 32% survival rate over the same time.
4. *Synthesis*. We found that high densities of turf algae can significantly inhibit recruitment of octocorals. Octocoral survival rates were similar to those published for scleractinians, but field settlement rates were much higher, which likely contributes to the higher resilience of octocorals to disturbances. The factors that influence recruitment are critical in understanding the dynamics of octocorals on Caribbean reefs as continuing declines in scleractinian cover may lead to more octocoral-dominated communities in the Caribbean.

## INTRODUCTION

Recruitment dynamics play a critical role in structuring and maintaining diversity in communities, and in the resilience of populations to disturbances (Warner and Chesson, 1985; Caley *et al.*, 1996; Wright, 2002). This is especially evident for sessile organisms, such as plants and benthic invertebrates, where variation in recruitment success is essential to recovery from disturbance and impacts adult population dynamics and spatial distributions and (Gaines and Roughgarden, 1985; McGuinness, 1996; Crawley, 2000; Price *et al.*, 2019). On Caribbean coral reefs the abundance of scleractinian corals has been decreasing for decades as a consequence of reduced recruitment in concert with bleaching events, hurricanes, and disease outbreaks (Goreau *et al.*, 1998; Gardner *et al.*, 2005; McWilliams *et al.*, 2005; Price *et al.*, 2019). Just as the decline in scleractinians is an inevitable consequence of failed recruitment, successful recruitment must have been essential to the increased prevalence of other taxa such as sponges and octocorals (Norström *et al.*, 2009; Ruzicka *et al.*, 2013; Lenz *et al.*, 2015; Edmunds and Lasker, 2016; Sánchez *et al.*, 2019).

Caribbean octocorals have always been a major constituent of contemporary Caribbean reef faunas (Kinzie, 1973). Coincident with the decline in scleractinian cover on some reefs, octocorals have increased in abundance (Ruzicka *et al.*, 2013; Lenz *et al.*, 2015; Edmunds and Lasker, 2016; Sánchez *et al.*, 2019). Octocorals have demonstrated greater resilience (ability to return to their previous population levels, *sensu* Gunderson, 2000) to disturbances than scleractinian corals (Edmunds and Lasker, 2016; Tsounis and Edmunds, 2017), but the basis of that resilience is still unknown. Reduced direct competition with scleractinians, higher fecundity, and greater juvenile survivorship and recruitment success have been suggested as potential reasons for their success in the Caribbean (Ruzicka *et al.*, 2013; Lenz *et al.*, 2015; Bartlett *et al.*, 2018). The lack of research concerning the mechanisms of resilience of octocorals is surprising, considering that octocoral forests are ecologically important (Kinzie, 1973; Williams *et al.*, 2015), provide some of the ecosystem functions that scleractinian reefs provide (e.g, Privitera-Johnson *et al.*, 2015; Tsounis *et al.*, 2016), and continue to maintain populations despite the increased prevalence of macroalgae.

Although recruitment is vital to maintaining coral populations (Yoshioka, 1996; Coles and Brown, 2007; McManus *et al.*, 2019), the interactions between benthic algae and octocoral recruitment are strikingly understudied, partly because of the difficulty in finding and repeatedly measuring newly-settled polyps (e.g., Linares *et al.*, 2007). The known effects of algae on the recruitment of octocorals are mixed; Alcolado *et al.* (2008) found no effect from macroalgae, while Linares *et al.* (2012) found a negative effect. Crusts from the family Corallinales promote settlement of octocorals (Lasker and Kim, 1996; Slattery *et al.*, 1999) while cyanobacteria may inhibit settlement (Kuffner *et al.*, 2006).

We differentiate between the two components of recruitment success: settlement and post-settlement survival. The distinction is important, as the two processes are often independent of each other. For example, while settlement patterns are critical to understanding the zonation of intertidal barnacles (Grosberg, 1982; Connell, 1985; Raimondi, 1988), post-settlement survival, not settlement, is the key factor in the establishment of distributional patterns in other systems (Grigg, 1988; Lasker *et al.*, 1998). Determining these differences has played an important role in understanding the ecology of recruitment and response to disturbance in many benthic taxa.

In this study, we used laboratory and field experiments to quantify the impacts of turf algae on the recruitment of newly-settled polyps of the octocoral *Plexaura homomalla* (Esper, 1792) by using tiles of differing levels of turf cover. Algal turf communities are a multispecies community of filamentous algae shorter than one centimeter (*sensu* Steneck and Dethier, 1994). They are highly diverse and productive and can have significant impacts on recruitment of sessile invertebrates (Adey and Steneck, 1985; Arnold *et al.*, 2010; Kramer *et al.*, 2012). We define recruitment as the addition of individuals to a population following settlement (*sensu* Caley *et al*., 1996). We hypothesize that recruitment will be negatively affected by algal turf cover. This work sheds light onto the interactions between newly-settled octocorals and the omnipresent turf algae that dominate Caribbean reefs (Hay, 1981; Birrell *et al.*, 2008). Understanding these processes will be important as further declines in scleractinian cover where they are still abundant could lead to more octocoral-dominated communities in the Caribbean (Tsounis and Edmunds, 2017).

## METHODS

### Collection, Spawning, and Larval Rearing

Female and male branches of *Plexaura homomalla* were collected from an octocoral-dominated reef in Round Bay, St. John, U.S. Virgin Islands (18.345 °N, 64.681 °W) on July 14-15, 2019 between 3.0 and 6.0 m depth. Colonies were determined to be gravid in the field by cutting a 5 cm piece diagonally and then looking for spermaries or eggs. Fragments of colonies were collected and transported to the Virgin Islands Environmental Resource Station and maintained in a sea table (i.e., flow-through, open-topped aquarium) with unfiltered running seawater pumped from Great Lameshur Bay (18.318 °N, 64.724 °W) from a depth of 1.5 m. Exchange rate in the sea table was approximately 200 L/h (60% of the volume hourly). Two EcoPlus 1/10 HP water chillers (Hawthorne Gardening Company, Vancouver, WA) were run in series to reduce the variability in temperature between day and night set to a temperature of 27°C. This reduced water temperature 0.5°C during the day (daily temperature range: 27.0-29.4°C) and maintained temperature at 27°C at night. Water temperature above 29°C is stressful to many Caribbean octocorals and can lead to bleaching (Prada *et al.*, 2010). Two submersible circulation pumps (Maxi-Jet 900 and Mini-Jet 404 [Marineland Spectrum Brands Pet LLC, Blacksburg, VA]) were placed within the tank to create a circular flow within the sea table.

Approximately two hours after sunset on the evenings of July 19-22, 2019 (3-6 days after full moon), both male and female *P. homomalla* colonies spawned in the tanks. Eggs from July 20 and 21 were used for the experiments. Pumps and water input were shut off at the start of spawning. Eggs were collected at least 30 minutes after spawning began to allow newly released eggs time to be fertilized. Eggs were collected and deposited into containers with 1 L of 10 µm filtered seawater (1500-2500 eggs/container). Shortly after collection, the gametes were diluted in 5 L polycarbonate containers with 3 L of 10 µm filtered seawater then reduced to 1 L through a 125 µm filter, which allowed the passage of sperm, but not eggs. This dilution was performed three times, leaving approximately 0.4% of the original seawater. This method reduces polyspermy, a potential source of early mortality in octocorals (Coelho and Lasker, 2016), and also removes decaying sperm that might promote bacterial growth in the cultures. During each of the subsequent four days, embryos were transferred to clean containers with minimal culture water to separate them from decaying gametes and embryos. These containers were filled to 5 L with 10 µm filtered seawater. After four days, embryos had developed into competent planulae.

### Laboratory Recruitment Experiment

Prior to the laboratory experiment, custom-fired stoneware clay tiles (n = 24, 14 × 14 × 1 cm) were deployed and conditioned on an octocoral-dominated reef in Grootpan Bay, St. John, U.S. Virgin Islands (18.309 °N, 64.719 °W) for 116 days. This site is also known as East Cabritte in other studies (e.g., Tsounis *et al.*, 2018; Lasker *et al.*, 2020). As refugia on tiles can improve recruitment rate of scleractinian larvae (Nozawa, 2008; Nozawa *et al.*, 2011), tiles were created with 0.5 cm deep and 1 cm wide pits and channels on one side. These substratum refugia were designed to examine the settlement preferences of octocoral and scleractinian larvae and post-settlement survival of octocoral larvae (Martínez-Quintana et al., in prep). Before the experiments, one of three treatments was applied to each tile (n = 8 per treatment): a control, herein referred to as “Reef”; scrubbed with a soft nylon bristle brush, herein referred to as “Scrubbed”; or removed from the reef and maintained in a sea table for 15 days, herein referred to as “Protected”. Reef tiles represented substrata subject to the natural grazing of fishes and invertebrates, Scrubbed tiles represented substrata that have been grazed more heavily, and Protected tiles represented substrata protected from grazing by macrofauna.

One day prior to introducing planulae, tiles were placed into 41 × 29 × 17 cm plastic containers (volume: 14 L, Sterilite Corporation, Townsend, MA). Containers were filled with 12 L of 10 µm filtered seawater. Two tiles from the same treatment were placed in each container on top of 1 cm square plastic mesh, rolled into 5 cm tall cylinders so that tiles were lifted off the bottom of the container to provide water circulation underneath. Pits and channels of the tiles were facing downwards. Water circulation in each container was provided by one glass Pasteur pipette attached to a Hydrofarm ActiveAQUA Air Pump Model PU110L (Hydrofarm LLC, Petaluma, CA). Water movement was slow but kept water moving throughout the container. Ambient sunlight was provided and temperature was held at 28°C.

Competent planulae (n = 150) were added to each container and allowed to settle for eight days and then polyps were counted. Planulae were competent when they readily attached to surfaces and were difficult to remove from the side of a glass pipette without forceful flushing of the water (Wells *et al.*, 2020). On day eight, no planulae were visible in the water and had either metamorphosed into polyps or died. 40% water changes were performed twice daily for the first two days and then daily afterwards, carefully checking for suspended planulae. Replacement seawater was 10-µm filtered for the first 6 days, after which seawater was 50-µm filtered. Motile invertebrates on the tiles were noted and identified to class or order. These tiles and associated polyps were used in the subsequent field survival and recruitment experiment.

### Field Recruitment Experiment

Nine days after adding planulae, the tiles were deployed onto the reef at Grootpan Bay, where the tiles were initially conditioned. Sampling was designed to span the early recruitment phase (first several weeks) and then resample at the end of two months. To do this, we counted the number of polyps on the undersides of the tiles on the day of deployment (day 0) and 2, 5, 9, 14, 19, and 57 days after deployment. Settlement of planulae from octocorals in the field occurred and often led to an increase in number of polyps over time, which confounded calculation of mortality rates of the original polyps. Polyps on nine of the 24 tiles (three from each treatment) were mapped on days 6, 9, 11, 14, 16, 19, and 57 in order to assess polyp survival and quantify settlement in the field. Distances between each polyp and its nearest-neighboring polyp were measured to the nearest millimeter for every individual on the nine mapped tiles to assess if settlement was aggregative, random, or dispersed.

Percent cover of living cover was determined using digital images of each tile taken on days 0, 19, and 57 taken with an Olympus Tough TG-5 or TG-6 12-megapixel waterproof digital camera (Olympus Corporation of the Americas, Center Valley, PA) and two Sola Dive Pro 2000 lights (Light & Motion, Marina, CA). Living cover groupings were broad to characterize large shifts in communities and were ascidians, bivalves, bryozoans, crustose coralline algae, chlorophytes, non-coralline crustose rhodophytes, sponges, turf algae, calcareous worm tubes, and the coral-overgrowing *Ramicrusta textilis* Pueschel & G.W. Saunders 2009. Images were analyzed with manual annotation using the online program CoralNet (Beijbom, 2015) with 100 randomly-generated points. This allowed us to quantify the differences in cover between the treatments and determine how the treatments changed after tiles were returned to the reef.

### Statistical Analyses

All statistical analyses were performed in R version 3.6.2 with the packages boot 1.3-23, coxme 2.2-16, lme4 1.1-21, survival 3.1-8, and vegan 2.5-6 (Bates *et al.*, 2015; Therneau, 2015; Canty and Ripley, 2017; Oksanen *et al.*, 2019; R Core Team, 2019; Therneau, 2020). Where confidence intervals were calculated, 95% bias-corrected and accelerated bootstrap confidence intervals with 10,000 replicates were used.

Effects of the algal cover treatments in the laboratory settlement experiment were analyzed using a generalized linear mixed effects model (GLMM) with a Poisson family distribution (Nelder and Wedderburn, 1972). The effects were modeled with treatment as a fixed effect and container as a random effect.

Analyses of polyp counts on the tiles after they were transferred to the reef can be divided into those based on settlement and survival on the nine mapped tiles and those based on the total number of polyps present on all 24 tiles. Survival data from the mapped tiles were analyzed based on the number of days each mapped polyp was alive related to algal turf cover. A mixed effect Cox proportional hazard model (CPH, Cox, 1972) was used to predict the effect of turf cover on survival of polyps on the mapped tiles. The model incorporated turf cover as a time-dependent and fixed covariate and tiles and polyps nested within tiles as random effects. CPH regression assumes that the relative effect of the hazard is proportional and consistent over time (Cox, 1972). Hence, the proportional hazards assumption was tested prior to the analysis. The analysis generates hazard ratios which depict the effects of varying turf cover on the probability of mortality relative to the probability of mortality at the median turf cover. Higher hazard ratios indicate a faster rate of mortality while a lower ratio indicates a reduced rate of mortality. Kaplan-Meier survival curves were used to visualize differences in polyp survival over time when exposed at different turf cover (i.e. turf covers of 0%, 25%, 50%, 75%, and 100%) from the hazard ratios calculated in the CPH.

Analyses of total numbers of polyps on the tiles after they were deployed in the field incorporates the effects of both mortality and field settlement. Field counts were first tested for normality with the Shapiro-Wilks W-test (Shapiro and Wilk, 1965). As data were not normal (Shapiro-Wilks, *p* < 0.05), counts were square root transformed and then a two-way repeated-measures analysis of variance (rANOVA) was performed with treatment and time as fixed factors and tile as a random factor.

Field settlement rates were calculated from the appearance of new polyps on mapped tiles divided by the total number of days observed. The day 19 to day 57 interval was excluded as all other time intervals were two or three days and mortality would obfuscate settlement during this longer time period. A nearest neighbor index (NN, Clark and Evans, 1954) was calculated for each tile to determine if settlement was aggregative, random, or evenly distributed in the field. The nearest-neighbor index equation was as follows:

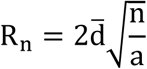

R_n_ is the nearest-neighbor statistic, 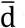 is the mean observed nearest-neighbor distance for the tile in cm, n is the number of polyps on the tile, and a is the area in cm^2^. An R_n_ of 1.0 is consistent with random settlement; values less than 1.0 suggest aggregative settlement and values greater than 1.0 suggest a more uniform pattern of settlement. This approach assumes that polyps cannot crawl across the benthos, which has not been described in octocoral settlers.

Living cover on the tiles at the end of the laboratory experiments and over the course of the field deployment was assessed using non-metric multidimensional scaling ordination plot (nMDS) based on Bray-Curtis dissimilarities between tiles.

## RESULTS

The amount and composition of cover on the tiles differed between treatments at the end of the laboratory experiment. Scrubbed tiles had low algal turf cover (0.1%), Reef tiles had middling turf cover (19%) and Protected tiles had high turf cover (50%). Turf was tallest in the Protected treatment (approximately 1 cm with some thalli almost 3.5 cm) and shortest in the Scrubbed treatment (no thalli above 0.5 cm). There were more motile invertebrates on the Protected tiles in the laboratory experiment, particularly amphipods, copepods, and small shrimp. Motile invertebrates were rarely observed on the Scrubbed tiles. Octocorals only recruited on tile surfaces. There was no recruitment on the chamber walls nor planulae in the water column at the end of the laboratory experiment. Recruitment was significantly different between tile treatments (GLMM, n = 12, *p* < 0.001). Protected tiles had low recruitment (11% of the initial planulae), Reef tiles had 38%, and Scrubbed tiles had 63% recruitment (Fig. 1).

**Fig. 1.**
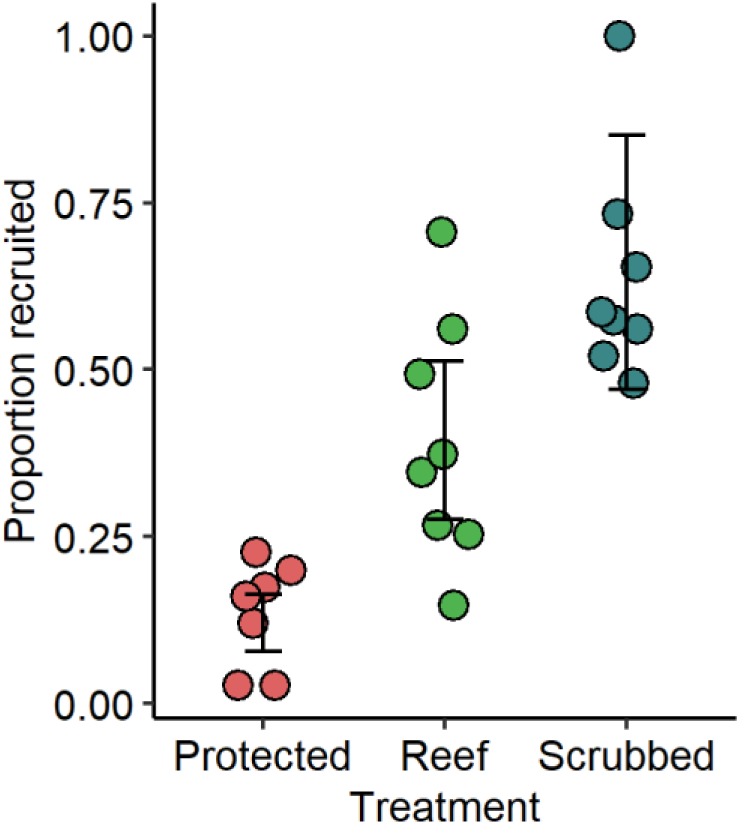
The effect of tile treatment on the proportion of octocoral polyps that recruited on the tiles in each container in the laboratory experiment. Treatments with reduced algal cover had greater numbers of recruits. Error bars are bootstrapped 95% confidence intervals of the means.

Once tiles were deployed in the field, the differences in the community structure due to treatment became less apparent over time (Supp. Fig. 1). On day 57, treatments were indistinguishable. This was primarily driven by bare space being replaced by turf. Observation of the tiles in the laboratory and field indicated that small crustaceans, which made up the bulk of the mesofaunal community, qualitatively declined after deployment. In the field, the only visible animals were gastropods, hermit crabs, and urchins. Copepods, amphipods, and annelids were rare on the tiles. Two bearded fireworms *Hermodice carunculata* (Pallas, 1766) were observed on tiles: one on a Protected tile and one on a Scrubbed tile. Observations of fireworms were concurrent with 58 and 72% reductions in octocoral abundance. Grazing by herbivorous fish on the sides and tops of the tiles was heavy and newly-settled polyps on those surfaces died quickly.

Analysis of the field data is complicated by two factors: settlement of naturally occurring planulae, which confounds estimates of mortality, and changes in the algal cover on the tiles, which affects the nature of the experimental treatments. When tiles were transferred to the field, they had a median of 21.5 polyps tile^-1^ (95% CI [11.5, 32.0]) and after 57 days had 5.0 polyps tile^-1^, (95% CI [3.0, 7.0]). However, counts on the tiles sometimes increased between observations due to settlement in the field. Multiple species appear to have contributed to the field settlement as the newly metamorphosed settlers had a range of colors and sizes. Therefore, survival rates of the polyps and the effects of turf on survival can only be determined for the polyps on the mapped tiles. Settlement and mortality on the mapped tiles are presented in Supp. Fig. 2. Among the mapped tiles, settlement was high, 8.7 polyps tile^-1^ 19 days^-1^ (95% CI [6.2, 11.3]), which can be extrapolated to 443 polyps m^-2^ 19 day^-1^ (95% CI [317, 577]) during this period of settlement. Positioning of new settlers on the tiles was random (NN, R_n_ = 1.05, 95% CI [0.94, 1.17], Supp. Fig. 3).

A total of 248 polyps were followed in assessing mortality rates on the mapped tiles. Mortality was 2.4% day^-1^, 95% CI (1.7%, 3.8%). Every 1% increase in algal turf cover was associated with a 1.3% increase in the mortality rate (CPH, *p* < 0.01, regression coefficient = 0.013, Fig. 2). When turf cover was modeled at 0%, 32% of octocoral polyps survived to 51 days. At modeled levels of 50% and 100% turf cover, survival dropped to 14% and 2%, respectively (Fig. 3). There was no relationship between turf cover and settlement rate on the mapped tiles (linear regression, *F*_1,41_ = 3.5, *p* = 0.66). Comparisons of the net change in numbers of polyps across all tiles identified a significant effect of treatment by time on the number of polyps (rANOVA, *F*_2,161_ = 3.5, *p =* 0.03, Fig. 4). Early in the experiment (day 0 to 5), Protected tiles had the same number or gained polyps, counts on Reef tiles stayed the same, and Scrubbed tiles lost polyps (Fig. 5A). Between days 5 and 14, mortality was offset by settlement on tiles across all treatments (Fig. 5B). By day 57, all but two tiles had lost polyps (Fig. 4) indicating that either settlement rates had declined and/or mortality had increased between days 19 and 57. Both tiles that gained polyps were initially Protected tiles. Changes in mortality rates in Scrubbed tiles disappeared when changes were evaluated as per capita rates. In that case, Reef and Scrubbed tiles had near-zero per capita rates (1.4% day^-1^, 95% CI [-1.2%, 8.1%] and −1.4% day^-1^, 95% CI [-4.4%, 0.8%], respectively). As settlement was random and therefore independent of density, and mortality was proportional to density, tiles with initially higher densities (Scrubbed tiles) were less likely to have settlement overcome mortality of polyps.

**Fig. 2.**
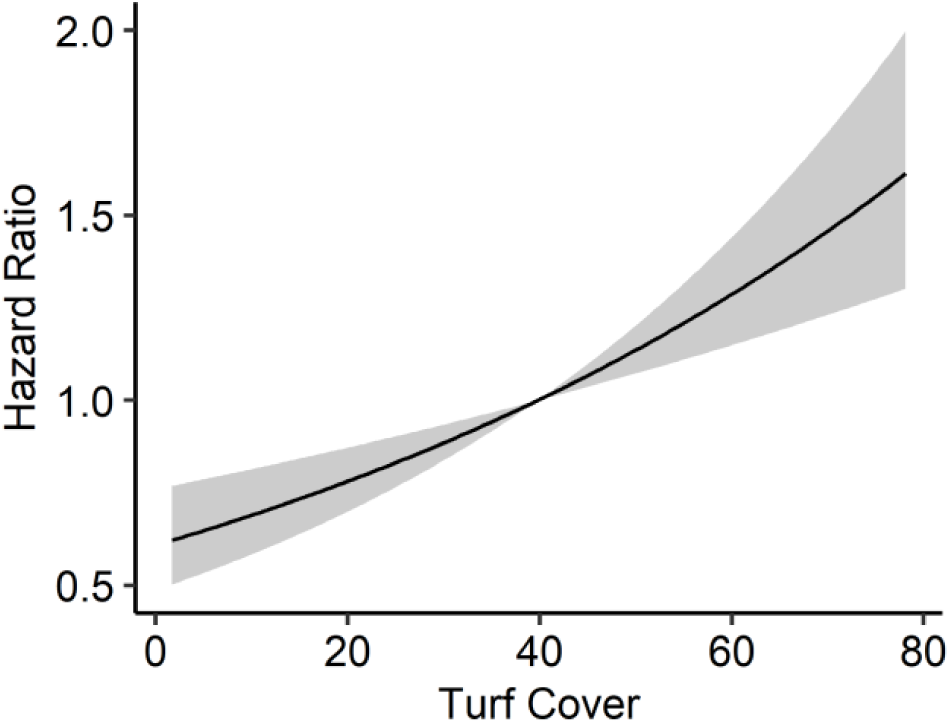
The relationship between algal turf cover and hazard ratio (i.e., ratio of the probability of mortality relative to the probability of mortality at the median turf cover [39%]). For every 1% increase in turf cover, octocorals polyps died 1.3% faster. The grey filled area is the 95% confidence interval.

**Fig. 3.**
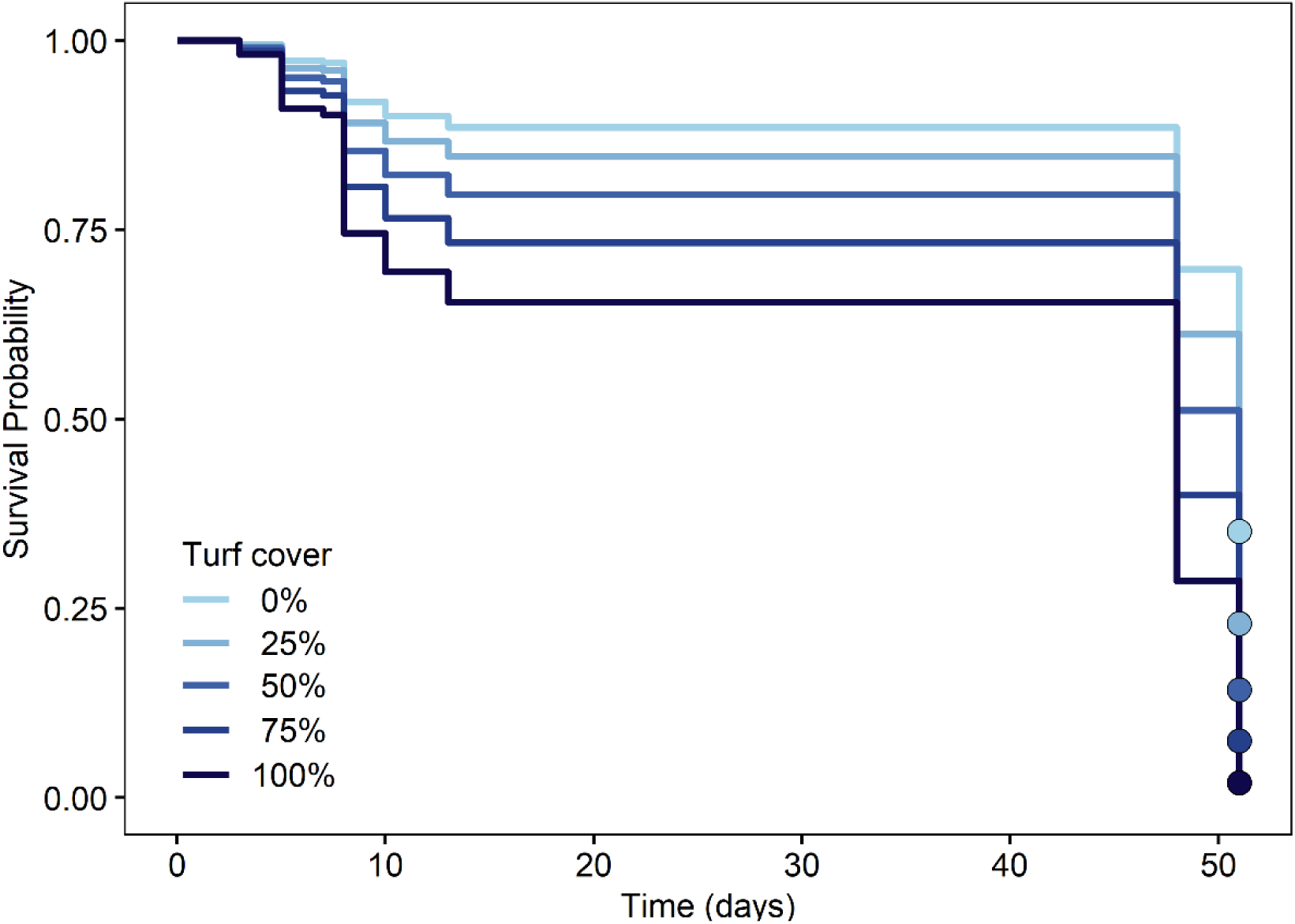
The modeled effect of constant algal turf cover on survival of octocoral polyps over time. Survival probability is significantly reduced with increased turf cover (Cox Mixed Effects Model, *p* = 0.03). The model does not interpolate survival between time points, creating the stepwise drops in survivorship. At the end of 51 days, survival probability was 16 times higher for individuals in 0% turf cover than in 100%. The endpoint of each model is indicated by a circle.

**Fig. 4.**
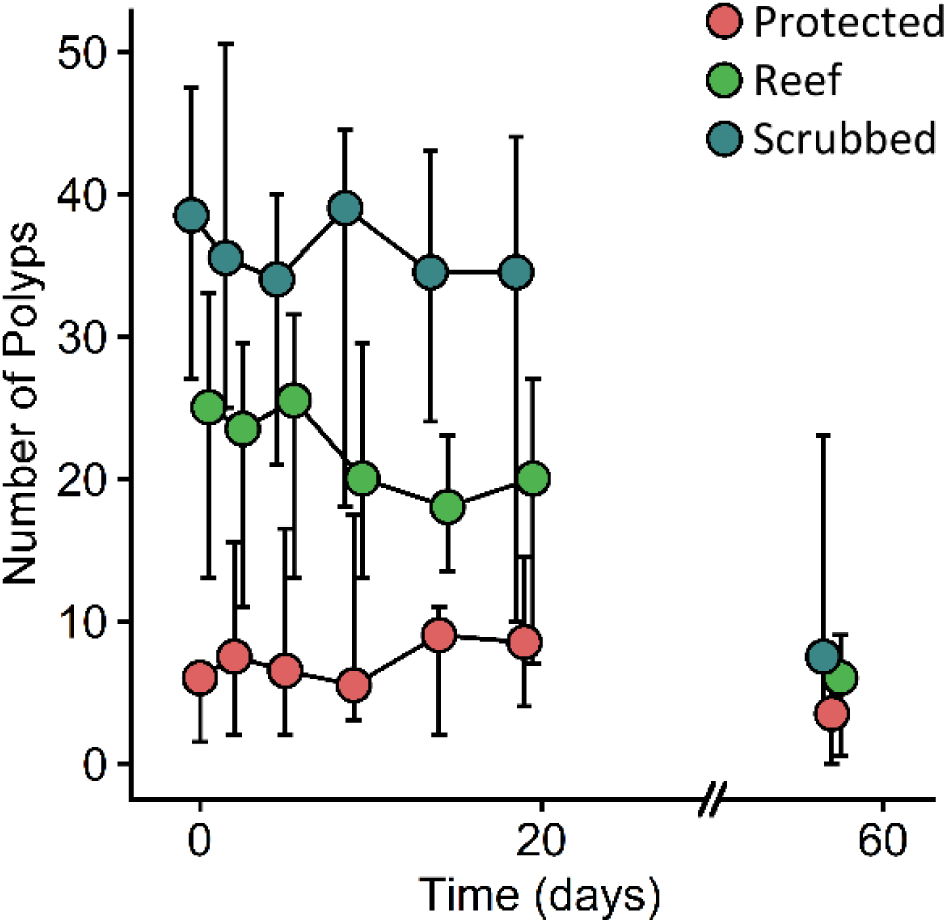
Change in median number of octocoral polyps over time by tile treatment. Scrubbed and Reef tiles decrease over time, while Protected tiles maintain a population of polyps. Data are median number of polyps. The last counts were performed on day 57. Error bars are bootstrapped 95% confidence intervals of the medians. Values have been offset on the x-axis to reduce overlap.

**Fig. 5.**
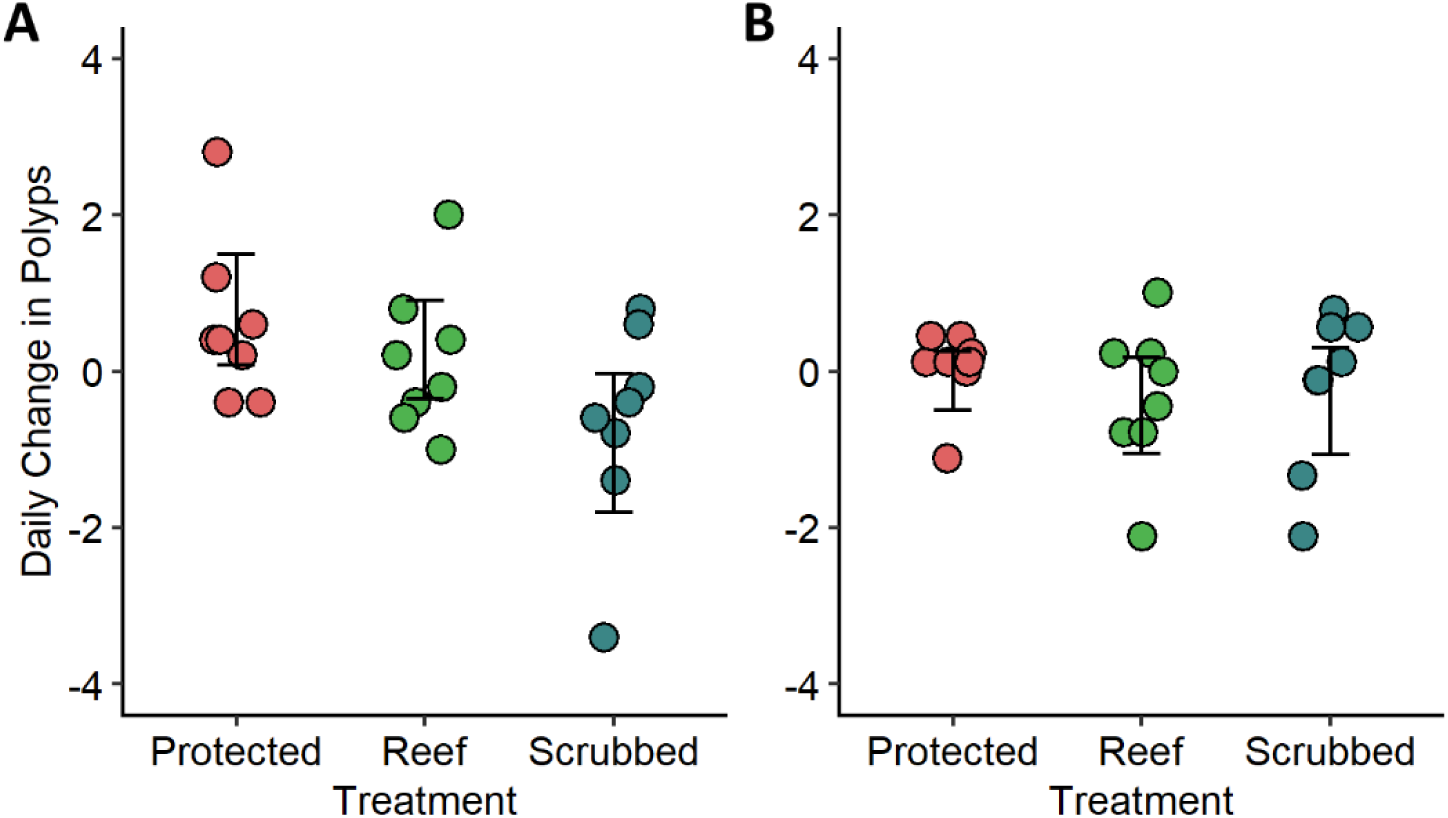
Daily change in number of polyps between (A) day 0 and day 5 and (B) day 5 and day 14. Early in the deployment, Protected tile mortality was overwhelmed by settlement of field-spawned octocorals, while Scrubbed tiles had lost polyps. Between days 5 and 14, mortality was offset by settlement on tiles across all treatments. Error bars are bootstrapped 95% confidence intervals of the means.

## DISCUSSION

While ecosystems naturally undergo changes in community composition, the prevalence and rates of change in modern communities are both high and accelerating (Dirzo *et al.*, 2014; Edmunds *et al.*, 2014; Ceballos *et al.*, 2015; Dornelas *et al.*, 2019). On Caribbean reefs, the past 4 decades have been witness to the precipitous decline in the abundance of scleractinian corals, one of the most abundant ecosystem engineers of the community (Gardner *et al.*, 2003; McWilliams *et al.*, 2005). At some sites these decadal declines have been stepwise, related to bleaching events and hurricanes, while other declines have been less episodic but no less dramatic (Gardner *et al.*, 2005). The decline in the abundance of sessile organisms, such as corals, is inevitably a consequence of the death of older individuals without replacement by new colonies (i.e. without successful recruitment). The decline in scleractinians has also been associated with dramatically greater bottom cover by algae (Hughes, 1994), and the effects of algal cover on recruitment have been cited as cause of the failed recovery of scleractinians (Birrell *et al.*, 2008). In contrast, the success of octocorals (Ruzicka *et al.*, 2013; Lenz *et al.*, 2015), including their recovery following disturbance events (Tsounis and Edmunds, 2017; Lasker *et al.*, 2020), suggests that recruitment of these species has continued regardless of the high cover of algae on most reefs. However, we discovered that octocorals, or at least *P. homomalla*, are not immune to the negative effects of algal turf.

High densities of turf algae significantly inhibited settlement and survival of juvenile octocorals. In the laboratory experiment there were striking differences in the number of planulae that successfully settled on the tiles in the different treatments (Fig. 1). As we did not observe settlement on the container walls, the differences in the numbers that settled reflects some combination of survival of planulae, successful settlement on the tile, and survival of polyps. In the field, higher algal cover was associated with increasing recruit mortality and had no relationship with settlement during the first two months following spawning (Fig 2).

Multiple hypotheses can account for the lower recruitment on tiles with higher algal cover. These include: 1) algae released toxic chemicals that affect local survival (i.e., allelopathy); 2) algae promoted the presence of pathogenic microbial populations; 3) algal and/or algal-associated bacterial respiration decreased oxygen to hypoxic or anoxic levels locally; 4) algal abrasion caused more frequent polyp retraction or damage; and 5) algae provided habitat for invertebrates that prey upon planulae and newly-settled polyps. These processes apply to settlement and survival in both our laboratory and field experiments. However, the relative magnitude of the effects in our laboratory setting was likely to have differed from field settings.

Allelopathy has been observed in interactions between algae and scleractinian corals, and has been proposed as a major barrier to scleractinian recovery following transitions to macroalgal dominance (Kuffner *et al.*, 2006; Rasher *et al.*, 2011; Bonaldo and Hay, 2014). Additionally, some algae can act as a reservoir for virulent scleractinian diseases (Nugues *et al.*, 2004; Casey *et al.*, 2014). In our experiments, allelopathic chemicals and/or pathogenic bacteria on the algae may have accumulated within the turf understory.

Algal turfs also have significant impacts on the diffusive boundary layer depending on canopy height and microscale substratum topography (Carpenter and Williams, 1993) and can drive oxygen levels down to hypoxic conditions at night (Brown and Carpenter, 2013). Extended anoxic and hypoxic conditions, such as what may be experienced by an octocoral polyp within turf algae, can lead to tissue necrosis in adult scleractinians (Smith *et al.*, 2006) and likely death in recruits. While flow was provided during the laboratory experiment and waves provided mixing in the field, algal respiration could have reduced oxygen to hypoxic levels within the diffusive boundary layer. In addition, algal turfs can also lead to hyperoxia during the day, which can stress the relationship between corals and their endosymbionts (reviewed in Lesser, 2011). This latter effect was unlikely in both our experiments as most recruits were on the underside of the tiles and light levels in the laboratory were low (indirect, ambient light from windows and ceiling mounted fluorescent lights).

Although it was an unlikely source of mortality in the laboratory, algal cover also leads to physical interactions between polyps and upright algae. These interactions can reduce growth by continually disturbing the colony, keeping the polyps closed for more time (River and Edmunds, 2001; Box and Mumby, 2007). When polyps are closed, colonies have less time to both capture prey and expose their endosymbionts to sunlight, and thus have lower growth. River and Edmunds (2001) found that abrasion-mediated polyp retraction led to an 80% reduction in growth rate. Flow rates were low within the settlement containers, but we frequently observed algae on tiles moving with the waves and this abrasion and disturbance could have increased mortality on the reef, especially in the denser turf mats.

An indirect and striking difference among the tiles in the laboratory experiments was the presence of mesofauna (amphipods, copepods, and small shrimps) in higher turf cover treatments. The abundance of mesofauna on the Protected treatment tiles was assumedly due to the elevated turf levels (50% cover vs. 19% and 0% in the other treatments) and the exclusion of grazing fishes and macroinvertebrates for 15 days prior to the start of the settlement experiment. The effect of mesofauna on the recruitment of octocorals is unknown, although some groups, such as cyclopoid and siphonostomatoid copepods, can be parasites of scleractinian corals (Cheng *et al.*, 2016) and can inhibit settlement of invertebrates through predation (Dahms *et al.*, 2004). Mesofaunal populations were lower in the field experiment, although shelled invertebrates (e.g., hermit crabs and snails), sea cucumbers, sea urchins, and worms were more abundant in the field, especially on the undersides of the tiles. On the two tiles with fireworms (Amphinomidae), polyp abundances were reduced by 58 and 72%. Fireworms are voracious predators of invertebrates including octocorals (Witman, 1988; Vreeland and Lasker, 1989) and likely had significant impacts in the field.

Despite the negative impacts of turf algae on octocoral recruitment, the survival rates in our experiments were greater than those reported for other Caribbean octocorals (Lasker *et al.*, 1998; Evans *et al.*, 2013). This was surprising given that we deployed younger polyps in the field, which are likely at a higher risk of mortality as octocorals generally follow a type III survival curve (Linares *et al.*, 2007). The higher survival in our experiments was likely due to refuge provided by the tiles. Both Lasker *et al.* (1998) and Evans *et al.* (2013) followed the fates of polyps on the upper side of settlement plates. In our study those polyps died quickly, likely through incidental predation by grazing herbivorous fish, and the field survivorship that we report was based on polyps on the underside of the tiles.

The combination of high settlement and low early mortality may be a major contributor to the high resilience to disturbance seen in octocorals, but not scleractinians. In this study, observed settlement rates for octocorals were higher than settlement rates of Caribbean scleractinian corals on tiles (e.g., Tomascik, 1991; Carlon, 2001; Green and Edmunds, 2011). For example, Green and Edmunds (2011) found an average scleractinian settlement rate of 76 polyps m^-2^ 6 months^-1^ on nearby reefs by counting skeletal remains on tiles after bleaching. In contrast, we observed settlement in the field more than five times as great in less than a month. Many octocoral species spawn for several months every year (Kahng *et al.*, 2011) and therefore settlement could be an order of magnitude greater than that of scleractinians. However, it is also important to recognize that recruitment rates may have been dramatically greater when scleractinians were far more abundant on Caribbean reefs (*cf.*, Sammarco, 1980, but also see Bak and Engel, 1979). Survival rates of octocorals fell within the range of survival rates published for scleractinians on tiles (Davies *et al.*, 2013; Trapon *et al.*, 2013; Doropoulos *et al.*, 2016). Davies *et al.* (2013) found that between 36 and 50% of *Agaricia* and *Porites* settlers survived the first 14 months, a higher survival rate than the 2 month survival of 33% that we observed for the octocoral polyps on our tiles. Doropoulos *et al.* (2016) found survival rates for scleractinian recruits between 13 and 47% after 50 days in the Pacific, and Trapon *et al.* (2013) found 42% survival after a month, similar to what we observed. Due to the difficultly of locating newly-settled corals on tiles, there are few studies looking at recruitment at the time scale of our observations. Simultaneous comparisons of octocoral and scleractinian settlement and survival are needed to determine if differences in recruitment can explain the differing trajectories of octocorals and scleractinians on many Caribbean Reefs. Future studies should include species-specific data with a particular emphasis on the survival within the first few months after settlement.

Our results suggest several directions where research should be productive. First, further research is required to understand the relationship between fish predators, mesofauna, and newly-settled octocorals. In this study, abundance of mesofauna seemed to be closely related to turf cover in the laboratory, suggesting that while fishes that prey on polyps will reduce survival, other fishes that consume mesofauna may have indirect positive effects on octocoral survival. Furthermore, the effects of the structural complexity of reef surfaces on survival must be further characterized. We observed differences in settlement and survival on the upper and lower surfaces of tiles. Lasker *et al.* (1998) reported differential settlement/survival of an octocoral on different structural elements of settlement plates and similar results have been reported for scleractinian recruitment rates (Edmunds *et al.*, 2014; Doropoulos *et al.*, 2016). However, the processes by which spatial complexity mediates facilitation, competition and/or predation is unknown. Finally, the effects of phenomena such as allelopathy, microbial load, and physiologically driven changes in water chemistry at the interface between the substratum and the water column are all poorly understood. Directed experiments looking at the effect of algal turfs on water quality within the benthic boundary layer may help explain distribution patterns of adult octocorals.

Caribbean reefs are undergoing a long-term increase in the abundance of octocorals while experiencing a reduction in scleractinian corals (Ruzicka *et al.*, 2013; Lenz *et al.*, 2015; Sánchez *et al.*, 2019). Those changes also affect the physical structure and ecosystem services provided by reef systems. In this study, we found that newly-settled octocorals do not survive as long in the presence of turfs, as has been reported for scleractinian corals (Birrell *et al.*, 2008; Penin *et al.*, 2011). However, on modern reefs octocorals may have greater settlement rates while maintaining a similar post-settlement survival, which suggests that octocorals might have higher early recruitment rates than scleractinians and that this process may be key in shaping the contemporary octocoral-dominated reef assemblages in the Caribbean.

## ACKNOWLEDGEMENTS

This work was completed under permits from the Virgin Islands National Park (VIIS-2019-SCI-0011) and the Virgin Islands Division of Fish and Wildlife (DFW19010) and was funded by the National Science Foundation (OCE 17-56381). We thank the Virgin Islands Environmental Resource Station (VIERS) and the University of the Virgin Islands (UVI) for laboratory space, L. Bramanti and A.M. Wilson for field and laboratory assistance, and E.R. Anderson and P.J. Edmunds for reading and commenting on the manuscript.

## REFERENCES

Adey, W.H. and Steneck, R.S. 1985. Highly productive eastern Caribbean reefs: Synergistic effects of biological, chemical, physical, and geological factors. Pages 163-187 in M. L. Reaka, editor. The Ecology of Coral Reefs. NOAA Undersea Research Program, Washington, DC, USA.

Alcolado, P.M., García-Parrado, P., Hernández-Muñoz, D. 2008. Estructura y composición de las comunidades de Gorgonias de los arrecifes del Archipiélago Sabana-Camagüey, Cuba: Conectividad y factores determinantes. Bulletin of Marine and Coastal Researcha 37:11–29.

Arnold, S.N., Steneck, R.S., Mumby, P.J. 2010. Running the gauntlet: Inhibitory effects of algal turfs on the processes of coral recruitment. Marine Ecology Progress Series 414:91–105.

Bak, R.P.M. and Engel, M.S. 1979. Distribution, abundance and survival of juvenile hermatypic corals (Scleractinia) and the importance of life history strategies in the parent coral community. Marine Biology 54:341–352.

Bartlett, L.A., Brinkhuis, V.I.P., Ruzicka, R.R., Collela, M.A., Semon Lunz, K., Leone, E.H., Hallock, P. 2018. Chapter 6. Dynamics of stony coral and octocoral juvenile assemblages following disturbance on patch reefs of the Florida Reef Tract. Pages 99-120 in C. Duque and Camacho, E. T., editors. Corals in a Changing World. IntechOpen, London, UK.

Bates, D., Mächler, M., Bolker, B., Walker, S. 2015. Fitting linear mixed-effects models using lme4. Journal of Statistical Software 67:48.

Beijbom, O. 2015. Automated annotation of coral reef survey images. Dissertation. University of California, San Diego, San Diego, CA, USA.

Birrell, C.L., McCook, L.J., Willis, B.L., Diaz-Pulido, G.A. 2008. Effects of benthic algae on the replenishment of corals and the implications for the resilience of coral reefs. Pages 25–63 in R. N. Gibson, Atkinson, R. J. A., and Gordon, J. D. M., editors. Oceanography and Marine Biology: An Annual Review. Taylor & Francis.

Bonaldo, R.M. and Hay, M.E. 2014. Seaweed-coral interactions: variance in seaweed allelopathy, coral susceptibility, and potential effects on coral resilience. PLoS ONE 9:e85786.

Box, S.J. and Mumby, P.J. 2007. Effect of macroalgal competition on growth and survival of juvenile Caribbean corals. Marine Ecology Progress Series 342:139–149.

Brown, A.L. and Carpenter, R.C. 2013. Water-flow mediated oxygen dynamics within massive *Porites*-algal turf interactions. Marine Ecology Progress Series 490:1–10.

Caley, M.J., Carr, M.H., Hixon, M.A., Hughes, T.P., Jones, G.P., Menge, B.A. 1996. Recruitment and the local dynamics of open marine populations. Annual Review of Ecology and Systematics 27:477–500.

Canty, A. and Ripley, B. 2017. boot: Bootstrap R (S-Plus) Functions.

Carlon, D.B. 2001. Depth-related patterns of coral recruitment and cryptic suspension-feeding invertebrates on Guana Island, British Virgin Islands. Bulletin of Marine Science 68:525–541.

Carpenter, R.C. and Williams, S.L. 1993. Effects of algal turf canopy height and microscale substratum topography on profiles of flow speed in a coral forereef environment. Limnology and Oceanography 38:687–694.

Casey, J.M., Ainsworth, T.D., Choat, J.H., Connolly, S.R. 2014. Farming behaviour of reef fishes increases the prevalence of coral disease associated microbes and black band disease. Proceedings of the Royal Society B: Biological Sciences 281.

Ceballos, G., Ehrlich, P.R., Barnosky, A.D., García, A., Pringle, R.M., Palmer, T.M. 2015. Accelerated modern human–induced species losses: Entering the sixth mass extinction. Science Advances 1:e1400253.

Cheng, Y.-R., Mayfield, A.B., Meng, P.-J., Dai, C.-F., Huys, R. 2016. Copepods associated with scleractinian corals: a worldwide checklist and a case study of their impact on the reef-building coral *Pocillopora damicornis* (Linnaeus, 1758) (Pocilloporidae). Zootaxa 4174:291–345.

Clark, P.J. and Evans, F.C. 1954. Distance to nearest neighbor as a measure of spatial relationships in populations. Ecology 35:445–453.

Coelho, M.A.G. and Lasker, H.R. 2016. Larval behavior and settlement dynamics of a ubiquitous Caribbean octocoral and its implications for dispersal. Marine Ecology Progress Series 561:109–121.

Coles, S.L. and Brown, E.K. 2007. Twenty-five years of change in coral coverage on a hurricane impacted reef in Hawai’i: The importance of recruitment. Coral Reefs 26:705–717.

Connell, J.H. 1985. The consequences of variation in initial settlement vs. post-settlement mortality in rocky intertidal communities. Journal of Experimental Marine Biology and Ecology 93:11–45.

Cox, D.R. 1972. Regression models and life tables. Journal of the Royal Statistical Society: Series B (Methodological) 34:187–220.

Crawley, M.J. 2000. Chapter 7: Seed predators and plant population dynamics. Pages 167–182 in M. Fenner, editor. Seeds: The Ecology of Regeneration in Plant Communities. CAB International.

Dahms, H.-U., Harder, T., Qian, P.-Y. 2004. Effect of meiofauna on macrofauna recruitment: settlement inhibition of the polychaete *Hydroides elegans* by the harpacticoid copepod *Tisbe japonica*. Journal of Experimental Marine Biology and Ecology 311:47–61.

Davies, S.W., Matz, M.V., Vize, P.D. 2013. Ecological complexity of coral recruitment processes: Effects of invertebrate herbivores on coral recruitment and growth depends upon substratum properties and coral species. PLoS ONE 8:e72830–e72830.

Dirzo, R., Young, H.S., Galetti, M., Ceballos, G., Isaac, N.J.B., Collen, B. 2014. Defaunation in the Anthropocene. Science 345:401–406.

Dornelas, M., Gotelli, N.J., Shimadzu, H., Moyes, F., Magurran, A.E., McGill, B.J. 2019. A balance of winners and losers in the Anthropocene. Ecology Letters 22:847–854.

Doropoulos, C., Roff, G., Bozec, Y.-M., Zupan, M., Werminghausen, J., Mumby, P.J. 2016. Characterizing the ecological trade-offs throughout the early ontogeny of coral recruitment. Ecological Monographs 86:20–44.

Edmunds, P.J. and Lasker, H.R. 2016. Cryptic regime shift in benthic community structure on shallow reefs in St. John, US Virgin Islands. Marine Ecology Progress Series 559:1–12.

Edmunds, P.J., Nozawa, Y., Villanueva, R.D. 2014. Refuges modulate coral recruitment in the Caribbean and the Pacific. Journal of Experimental Marine Biology and Ecology 454:78–84.

Evans, M.J., Coffroth, M.A., Lasker, H.R. 2013. Effects of predator exclusion on recruit survivorship in an octocoral (*Briareum asbestinum*) and a scleractinian coral (*Porites astreoides*). Coral Reefs 32:597–601.

Gaines, S.D. and Roughgarden, J. 1985. Larval settlement rate: A leading determinant of structure in an ecological community of the marine intertidal zone. Proceedings of the National Academy of Sciences 82:3707.

Gardner, T.A., Côté, I.M., Gill, J.A., Grant, A., Watkinson, A.R. 2003. Long-term region-wide declines in Caribbean corals. Science 301:958–960.

Gardner, T.A., Côté, I.M., Gill, J.A., Grant, A., Watkinson, A.R. 2005. Hurricanes and Caribbean coral reefs: Impacts, recovery patterns, and role in long-term decline. Ecology 86:174–184.

Goreau, T.F., Cervino, J., Goreau, M., Hayes, R., Hayes, M., Richardson, L., Smith, G., DeMeyer, K., Nagelkerken, I., Garzon-Ferrera, J., Gil, D., Garrison, G., Williams, E.H., Bunckley-Williams, L., Quirolo, C., Patterson, K., Porter, J.W., Porter, K. 1998. Rapid spread of diseases in Caribbean coral reefs. Revista de Biología Tropical 46:157–171.

Green, D.H. and Edmunds, P.J. 2011. Spatio-temporal variability of coral recruitment on shallow reefs in St. John, US Virgin Islands. Journal of Experimental Marine Biology and Ecology 397:220–229.

Grigg, R.W. 1988. Recruitment limitation of a deep benthic hard-bottom octocoral population in the Hawaiian Islands. Marine Ecology Progress Series 45:121–126.

Grosberg, R.K. 1982. Intertidal zonation of barnacles: The influence of planktonic zonation of larvae on vertical distribution of adults. Ecology 63:894–899.

Gunderson, L.H. 2000. Ecological resilience - in theory and application. Annual Review of Ecology and Systematics 31:425–439.

Hay, M.E. 1981. The functional morphology of turf-forming seaweeds: persistence in stressful marine habitats. Ecology 62:739–750.

Hughes, T.P. 1994. Catastrophes, phase shifts, and large-scale degradation of a Caribbean coral reef. Science 265:1547–1551.

Kahng, S.E., Benayahu, Y., Lasker, H.R. 2011. Sexual reproduction in octocorals. Marine Ecology Progress Series 443:265–283.

Kinzie, R.A., III. 1973. The zonation of West Indian gorgonians. Bulletin of Marine Science 23:93–155.

Kramer, M.J., Bellwood, D.R., Bellwood, O. 2012. Cryptofauna of the epilithic algal matrix on an inshore coral reef, Great Barrier Reef. Coral Reefs 31:1007–1015.

Kuffner, I.B., Walters, L.J., Becerro, M.A., Paul, V.J., Ritson-Williams, R., Beach, K.S. 2006. Inhibition of coral recruitment by macroalgae and cyanobacteria. Marine Ecology Progress Series 323:107–117.

Lasker, H.R., Kiho, K., Coffroth, M.A. 1998. Production, settlement, and survival of plexaurid gorgonian recruits. Marine Ecology Progress Series 162:111–123.

Lasker, H.R. and Kim, K. 1996. Larval development and settlement behavior of the gorgonian coral *Plexaura kuna* (Lasker, Kim and Coffroth). Journal of Experimental Marine Biology and Ecology 207:161–175.

Lasker, H.R., Martínez-Quintana, Á., Bramanti, L., Edmunds, P.J. 2020. Resilience of octocoral forests to catastrophic storms. Scientific Reports 10:4286.

Lenz, E.A., Bramanti, L., Lasker, H.R., Edmunds, P.J. 2015. Long-term variation of octocoral populations in St. John, US Virgin Islands. Coral Reefs 34:1099–1109.

Lesser, M.P. 2011. Coral bleaching: Causes and mechanisms. Pages 405–419 in Z. Dubinsky and Stambler, N., editors. Coral Reefs: An Ecosystem in Transition. Springer Netherlands, Dordrecht.

Linares, C., Cebrian, E., Coma, R. 2012. Effects of turf algae on recruitment and juvenile survival of gorgonian corals. Marine Ecology Progress Series 452:81–88.

Linares, C., Doak, D.F., Coma, R., Díaz, D., Zabala, M. 2007. Life history and viability of a long-lived marine invertebrate: The octocoral *Paramuricea clavata*. Ecology 88:918–928.

McGuinness, K.A. 1996. Dispersal, establishment and survival of *Ceriops tagal* propagules in a north Australian mangrove forest. Oecologia 109:80–87.

McManus, L.C., Watson, J.R., Vasconcelos, V.V., Levin, S.A. 2019. Stability and recovery of coral-algae systems: the importance of recruitment seasonality and grazing influence. Theoretical Ecology 12:61–72.

McWilliams, J.P., Côté, I.M., Gill, J.A., Sutherland, W.J., Watkinson, A.R. 2005. Accelerating impacts of temperature-induced coral bleaching in the Caribbean. Ecology 86:2055–2060.

Nelder, J.A. and Wedderburn, R.W.M. 1972. Generalized linear models. Journal of the Royal Statistical Society: Series A (General) 135:370–384.

Norström, A.V., Nyström, M., Lokrantz, J., Folke, C. 2009. Alternative states on coral reefs: beyond coral–macroalgal phase shifts. Marine Ecology Progress Series 376:295–306.

Nozawa, Y. 2008. Micro-crevice structure enhances coral spat survivorship. Journal of Experimental Marine Biology and Ecology 367:127–130.

Nozawa, Y., Tanaka, K., Reimer, J.D. 2011. Reconsideration of the surface structure of settlement plates used in coral recruitment studies. Zoological Studies 50:53–60.

Nugues, M.M., Smith, G.W., van Hooidonk, R.J., Seabra, M.I., Bak, R.P.M. 2004. Algal contact as a trigger for coral disease. Ecology Letters 7:919–923.

Oksanen, J., Blanchet, F.G., Friendly, M., Kindt, R., Legendre, P., McGlinn, D., Minchin, P.R., O’Hara, R.B., Simpson, G.L., Solymos, P., Stevens, H.H., Szoecs, E., Wagner, H. 2019. vegan: Community Ecology Package.

Penin, L., Michonneau, F., Carroll, A., Adjeroud, M. 2011. Effects of predators and grazers exclusion on early post-settlement coral mortality. Hydrobiologia 663:259–264.

Prada, C., Weil, E., Yoshioka, P.M. 2010. Octocoral bleaching during unusual thermal stress. Coral Reefs 29:41–45.

Price, N.N., Muko, S., Legendre, L., Steneck, R.S., van Oppen, M.J.H., Albright, R., Ang, P., Jr., Carpenter, R.C., Chui, A.P.Y., Fan, T.Y., Gates, R.D., Harii, S., Kitano, H., Kurihara, H., Mitarai, S., Padilla-Gamiño, J.L., Sakai, K., Suzuki, G., Edmunds, P.J. 2019. Global biogeography of coral recruitment: Tropical decline and subtropical increase. Marine Ecology Progress Series 621:1–17.

Privitera-Johnson, K., Lenz, E.A., Edmunds, P.J. 2015. Density-associated recruitment in octocoral communities in St. John, US Virgin Islands. Journal of Experimental Marine Biology and Ecology 473:103–109.

R Core Team. 2019. R: A Language and Environment for Statistical Computing in R Foundation for Statistical Computing, editor. R Foundation for Statistical Computing, Vienna, AT.

Raimondi, P.T. 1988. Rock type affects settlement, recruitment, and zonation of the barnacle *Chthamalus anisopoma* Pilsbury. Journal of Experimental Marine Biology and Ecology 123:253–267.

Rasher, D.B., Stout, E.P., Engel, S., Kubanek, J., Hay, M.E. 2011. Macroalgal terpenes function as allelopathic agents against reef corals. Proceedings of the National Academy of Sciences 108:17726.

River, G.F. and Edmunds, P.J. 2001. Mechanisms of interaction between macroalgae and scleractinians on a coral reef in Jamaica. Journal of Experimental Marine Biology and Ecology 261:159–172.

Ruzicka, R.R., Colella, M.A., Porter, J.W., Morrison, J.M., Kidney, J.A., Brinkhuis, V., Lunz, K.S., Macaulay, K.A., Bartlett, L.A., Meyers, M.K., Colee, J. 2013. Temporal changes in benthic assemblages on Florida Keys reefs 11 years after the 1997/1998 El Niño. Marine Ecology Progress Series 489:125–141.

Sammarco, P.W. 1980. *Diadema* and its relationship to coral spat mortality: Grazing, competition, and biological disturbance. Journal of Experimental Marine Biology and Ecology 45:245–272.

Sánchez, J.A., Gómez-Corrales, M., Gutierrez-Cala, L., Vergara, D.C., Roa, P., González-Zapata, F.L., Gnecco, M., Puerto, N., Neira, L., Sarmiento, A. 2019. Steady decline of corals and other benthic organisms in the SeaFlower Biosphere Reserve (Southwestern Caribbean). Frontiers in Marine Science 6:73.

Shapiro, S.S. and Wilk, M.B. 1965. An analysis of variance test for normality (complete samples). Biometrika 52:591–611.

Slattery, M., Hines, G.A., Starmer, J., Paul, V.J. 1999. Chemical signals in gametogenesis, spawning, and larval settlement and defense of the soft coral *Sinularia polydactyla*. Coral Reefs 18:75–84.

Smith, J.E., Shaw, M., Edwards, R.A., Obura, D., Pantos, O., Sala, E., Sandin, S.A., Smriga, S., Hatay, M., Rohwer, F.L. 2006. Indirect effects of algae on coral: Algae-mediated, microbe-induced coral mortality. Ecology Letters 9:835–845.

Steneck, R.S. and Dethier, M.N. 1994. A functional group approach to the structure of algal-dominated communities. Oikos 69:476–498.

Therneau, T.M. 2015. A package for survival analysis in S.

Therneau, T.M. 2020. coxme: Mixed Effects Cox Models.

Tomascik, T. 1991. Settlement patterns of Caribbean scleractinian corals on artificial substrata along a eutrophication gradient, Barbados, West Indies. Marine Ecology Progress Series 77:261–269.

Trapon, M.L., Pratchett, M.S., Hoey, A.S., Baird, A.H. 2013. Influence of fish grazing and sedimentation on the early post-settlement survival of the tabular coral *Acropora cytherea*. Coral Reefs 32:1051–1059.

Tsounis, G. and Edmunds, P.J. 2017. Three decades of coral reef community dynamics in St. John, USVI: A contrast of scleractinians and octocorals. Ecosphere 8:e01646.

Tsounis, G., Edmunds, P.J., Bramanti, L., Gambrel, B., Lasker, H.R. 2018. Variability of size structure and species composition in Caribbean octocoral communities under contrasting environmental conditions. Marine Biology 165:29.

Tsounis, G., Steele, M., Edmunds, P.J. 2016. Effects of octocorals on the feeding behavior of herbivorous coral reef fishes in the Caribbean. Pages 118–119 in Western Society of Naturalists, Monterey, CA, USA.

Vreeland, H.V. and Lasker, H.R. 1989. Selective feeding of the polychaete *Hermodice carunculata* Pallas on Caribbean gorgonians. Journal of Experimental Marine Biology and Ecology 129:265–277.

Warner, R.R. and Chesson, P.L. 1985. Coexistence mediated by recruitment fluctuations: A field guide to the storage effect. The American Naturalist 125:769–787.

Wells, C.D., Tonra, K.J., Lasker, H.R. 2020. Embryogenesis, polyembryony, and settlement in the gorgonian *Plexaura homomalla*. bioRxiv:2020.2003.2019.999300.

Williams, S.M., Chollett, I., Roff, G., Cortés, J., Dryden, C.S., Mumby, P.J. 2015. Hierarchical spatial patterns in Caribbean reef benthic assemblages. Journal of Biogeography 42:1327–1335.

Witman, J.D. 1988. Effects of predation by the fireworm *Hermodice carunculata* on milleporid hydrocorals. Bulletin of Marine Science 42:446–458.

Wright, J.S. 2002. Plant diversity in tropical forests: a review of mechanisms of species coexistence. Oecologia 130:1–14.

Yoshioka, P.M. 1996. Variable recruitment and its effects on the population and community structure of shallow-water gorgonians. Bulletin of Marine Science 59:433–443.

