## Supplemental Figures 1-3 for "Welcome to the jungle: Algal turf negatively affects recruitment of a Caribbean octocoral"

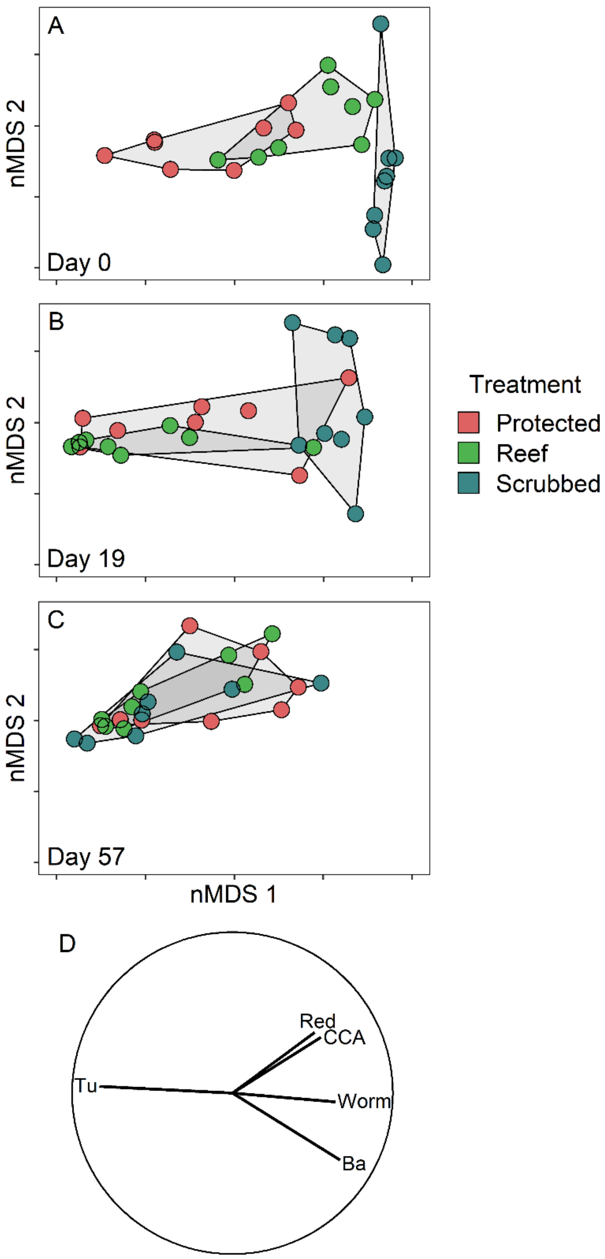


Supp. Fig. 1 Non-metric multidimensional scaling plot of the change in the multivariate community structure for the tile communities (A) at the end of the laboratory experiment and start of the field experiment (day 0), (B) on day 19, and (C) day 57. Plots were produced from the same ordination (stress = 0.09), but were plotted seperately. (D) Vectors display Spearman correlations between taxa and nMDS 1 and nMDS 2 (only values above 0.6 displayed). Through time, treatments became more similar, becoming indistinguishable. Separation along nMDS 1 was primarily associated with abundance of turf algae and calcified worm tubes. Separation along nMDS 2 was associated with algal crusts and bare surface. Ba: bare surface; CCA: crustose coralline algae; Red: non-CCA red algal crusts; Tu: turf; Worm: calcified worm tubes.

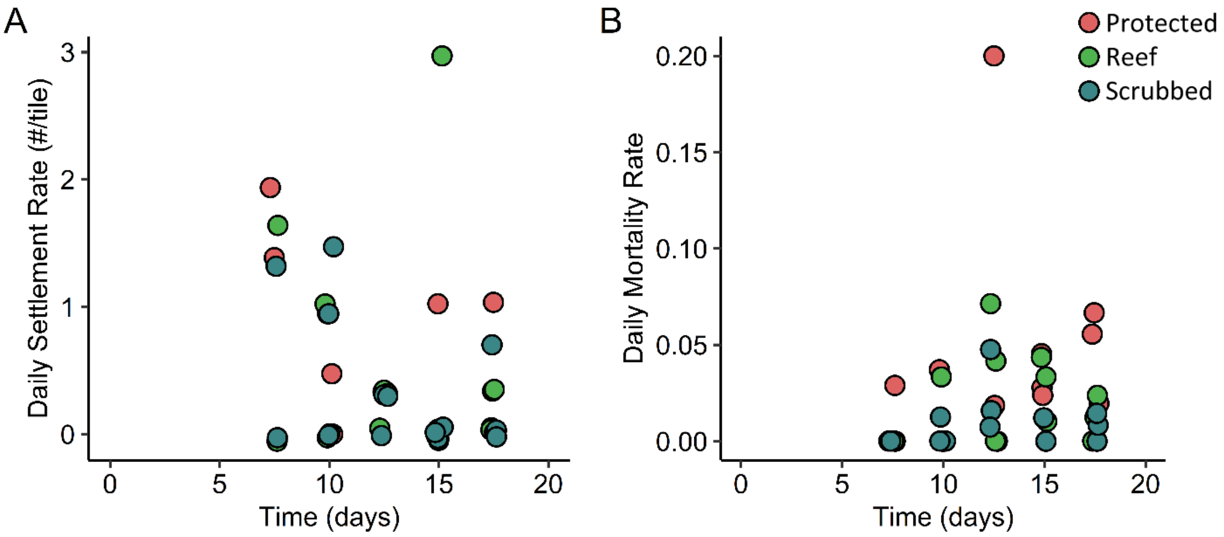


Supp. Fig. 2. Daily (A) settlement and (B) mortality rate of octocoral polyps on the mapped tiles.

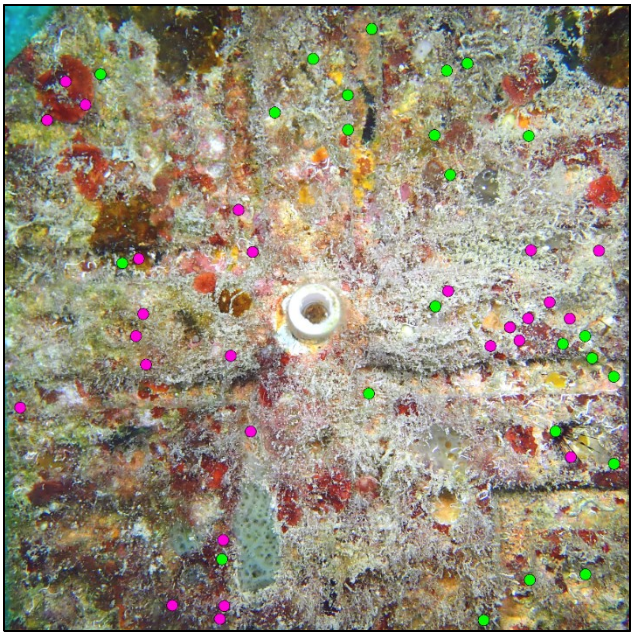

Supp. Fig. 3. A mapped tile illustrating the location of all planulae settlement locations on day 57. Green-filled circles are locations that had live polyps on day 57 and pink-filled circles are locations where polyps had died by day 57. Settlement position of polyps in the field was random ($R_n$ = 1.05, 95% CI [0.94, 1.17]).
